# Pulsed Electromagnetic Fields Alleviates Hepatic Oxidative Stress and Lipids Accumulation in db/db mice

**DOI:** 10.1101/2020.04.06.028621

**Authors:** Ying Liu, Mingming Zhai

**Author notes:** Corresponding Author: Mingming Zhai, Department of Biomedical Engineering, Fourth Military Medical University, 169 Changle West Road, Xi’an, China. First Author: Ying Liu, EMAIL (Ying Liu.).

## Abstract

**Purpose:** Nonalcoholic fatty liver disease (NAFLD), affected more than 70 % of patients with type 2 diabetes (T2DM), has become a common metabolic liver disease worldwide. However, the specifically treatments targeting NAFLD have not been found until now. Pulsed electromagnetic fields have positive effects on multiple diseases. However, the effects of PEMF on NAFLD in T2DM require further investigation. The present study assessed the effects of pulsed electromagnetic fields on the liver oxidative stress and lipid accumulation of db/db mice.

**Patients and methods:** Animals were exposed to 2 h of pulsed electromagnetic fields (15.38 Hz, 2 mT) or sham stimulated, and thereafter sacrificed at 8 weeks later. The biomarkers of oxidative stress, such as MDA, GSSG and GSH levels, were analysed with commercial kits. The activity of liver antioxidant enzymes as CAT, SOD and GSH-Px was detected. Hepatic expressions of CAT, GR, GSH-Px, SOD1, SOD2 and SREBP-1c at protein levels were determined with Western blotting. Hepatic weight was measured and triglyceride accumulation were visualized by Oil Red O staining.

**Results:** PEMF exposure could protect the liver from oxidative stress injury by decreasing MDA and GSSG level, promoting reduced GSH level, and increasing GSH-Px activity and expression in comparison with sham group. But CAT and SOD activity have no statistic difference as same as CAT, GR, SOD1 and SOD2 expression. Furthermore, PEMF exposure reduced liver weight and triglyceride content. Meanwhile, PEMF exposure ameliorated hepatic steatosis through reducing the expression of SREBP-1c to regulate the lipid synthesis.

**Conclusion:** The present study provides evidence that PEMF could increase antioxidant enzymes activity and alleviate lipid accumulation in fatty liver. This implies that PEMF exposure has beneficial effects for the treatment of NAFLD in accompany with T2DM.

## Introduction

Type 2 diabetes mellitus (T2DM) is the most common form of diabetes with metabolic disturbance. Recently literatures identified that almost 70% of T2DM patients present with non-alcoholic fatty liver disease (NAFLD) and the pathophysiology of NAFLD is still under illumination^1, 2^. A closely connection between NAFLD and T2DM has increasingly been identified. Notably, a large number of patients with NAFLD is associated with an increased risk of incident T2DM^3, 4^. But to date, the specifical drug therapy approved for NAFLD have not been found^5, 6^.

NAFLD is not caused by alcohol consumption, but associated with hepatic steatosis or accumulation of lipids, predominantly triglycerides, within hepatocytes. Although the pathophysiological mechanism with underlying NAFLD has not being elucidated yet, researches have suggested that oxidative stress is one of the main factors effecting insulin resistance and the subsequent progression of NAFLD. Excessive generation of reactive oxygen species (ROS) along with reduced activity of antioxidant enzymatic would interfere with insulin signalling negatively which cause insulin resistance and increased triglyceride (TG) accumulation in the liver^7^. Liver is Serving as a hub of lipid metabolism, has abundant of mitochondria, and thus vulnerable to ROS attack. Generally, liver functions as a major target of a variety of hormones to regulate glycogenolysis and gluconeogenesis. Insulin as the only hormone plays an important role with lowering blood glucose. Therefore, the liver responds to insulin abnormally that would lead to the regulation of blood glucose out of control. Thus, ameliorating liver oxidative stress is hypothesized to be one important strategy for treatment of NAFLD patients with T2DM. Clinical studies have attempted to treat diabetic complications with synthetic antioxidant therapy. However, the effectiveness of these results were not as expected^8, 9^. In recent years, physical factor therapies improved oxidative stress have been proved^10-13^. Especially, the pulsed electromagnetic fields (PEMF) as a safe, non-invasive treatment method is approved by U.S. Food and Drug Administration (FDA)^14^. Some evidences have found that PEMF as an alternative method are capable to improve oxidative stress. PEMF stimulation can suppress the production of ROS in human osteoblasts, indicating alterations in cells antioxidative stress response^12^. Additionally, PEMF exposure reduced the ROS-level and upregulated the expression of catalase (CAT) and superoxide dismutase (SOD) in rats, suggesting modulations in oxidative stress accompany with spinal cord injury ^11^.

Thus, this present study aims to investigate whether the PEMF exposure has beneficial effects on hepatic oxidative stress and lipid accumulation in db/db mice, a model of T2DM widely utilized to study NAFLD, and to further explore the underlining mechanisms. The results indicated that PEMF could ameliorate oxidative stress and alleviate hepatic steatosis in db/db mice.

## Methods and materials

### Materials and Reagents

BCA Protein Assay Kit was purchased from Thermo Scientific. SREBP-1c antibody was purchased from Santa Cruz Biotechnology. CAT antibody was purchased from Proteintech Group. Glutathione reductase (GR), β-Actin and glutathione peroxidase (GSH-Px) antibodies were purchased from Bioworld Technology. SOD1 and SOD2 antibodies were purchased from Abcam Biotechonogy. The kits used to test the concentration of hepatic TG, MDA, GSH and GSSG, and the activity of hepatic CAT, SOD and GSH-Px were purchased from Nanjing Jiancheng Bioengineering Institute. Other chemicals used in this study were purchased from Sigma.

### Animals and Experimental Procedures

Animal experiments were approved by Fourth Military Medical University Ethical committee. Eight 8-week-old female C57BL/KsJ mice (17.87±1.33 g) and 16 8-week-old female C57BL/KsJ db/db mice (29.51±2.37 g), originally obtained from Jackson Laboratory, were purchased from Model Animal Research Centre of Nanjing University where they were bred. The db/db mice were raised in a standard environmental condition with 12-h light/12-h dark cycle and temperature control (22±1 °C). They were free to consume standard chow and water.

The female C57BL/KsJ mice set as control group (n=8), and the 16 female C57L/6J db/db mice were randomly divided into two groups including sham group (n=8) and PEMF group (n=8). All the mice of PEMF group were exposed for 12 weeks and 2 h/day with 2 mT. The control and sham group were placed in an identical chamber with no pulsed electromagnetic fields. At the end of the experiment, the mice were anesthetized by gas and hepatic samples were harvested for the assays. Hepatic samples used for determination of MDA, reduced glutathione (GSH) and oxidized GSH (GSSG), analysis of antioxidation enzymes activity, western blotting, and TG content determination were stored at −80 °C until use. Hepatic frozen sections for lipid accumulation analysis were sliced immediately.

### Electromagnetic Exposure System

PEMF stimulators in this study were described previously (GHY-III, FMMU, Xi’an, China; China Patent No. ZL02224739.4)^15-17^. In brief, the waveform consist of a pulse burst (burst width, 5 ms; pulse wait, 0.02 ms; burst wait, 60 ms; pulse rise, 0.3 μs; pulse fall, 2.0 μs) repeated at 15.38 Hz (Fig. 1). Interval distance between two coils (20 cm diameter) were 10 cm, and turn numbers of enamel-coated round copper wire (1.0 mm diameter) was 80. To monitor waveform and amplitude of the current in the coils, a high-power resistor of 1 Ω was placed in series with the coils and voltage drop across the resistor was observed with an oscilloscope (Agilent Technologies, Santa Clara, CA) during PEMF exposure period. In this study, the baseline level of magnetic field was 0 mT, and determined peak magnetic field intensity of coils was approximately 2 mT. Accuracy for peak magnetic field measurement was further confirmed using a Gaussmeter (Model 455 DSP, Lake Shore Cryotronics, Westerville, OH) with a transversal Hall Probe (HMFT-3E03-VF). The measured background electromagnetic field was 50±2 μT.

**Figure 1:**
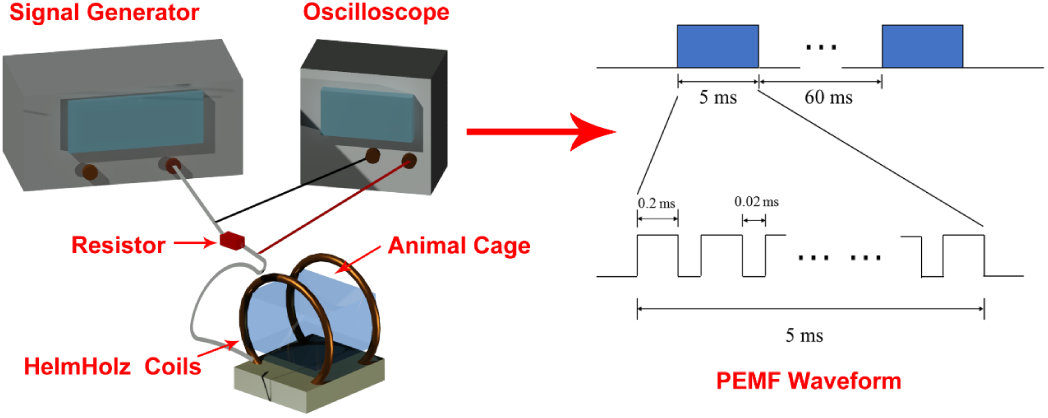
Schematic representation of PEMF exposure system and output waveform. The exposure waveform comprises a pulsed burst (burst width, 5 ms; pulse width, 0.2 ms; pulse wait, 0.02 ms; burst wait, 60 ms; pulse rise, 0.3 μs; pulse fall, 2.0 μs) repeated at 15.38 Hz.

### Biochemical Assays for Hepatic MDA and GSH Content and Antioxidation Enzymes

To measure MDA, GSH and GSSG level as well as the activity of antioxidation enzymes in livers, commercial kits were used. Briefly, to determine MDA level and activities of antioxidant enzymes, the liver samples were homogenized in the ice-water bath with 0.9 % saline at the rate of 100 mg : 0.9 ml. Next, the homogenate was centrifuged at 570 g for 10 min. The supernatant was collected and determined under the instructions. For testing GSH and GSSG level, the liver samples were homogenized in ice-water bath with agent given in the kit at the rate of 200 mg : 0.8 ml. Then, the homogenate was centrifuged at 1120 g for 10 min. The supernatant was collected and measured in accordance with the instructions.

### Oil Red O Staining

To explore the lipid accumulation in the hepatic cells, frozen liver samples were sliced at 10 μm and stained with Oil Red O solution. The frozen slices were placed into the working solution and stained for 15 min at 22 °C. Then, the slices were washed with 37 °C distilled water for 15 s. At last, the sections were observed with an Olympus light microscope.

### Biochemical Assay for Hepatic Triglyceride

To determine the hepatic triglyceride, a commercial kit was taken. In brief, the samples were homogenized with absolute ethyl alcohol at the rate of 100 mg : 0.9 ml in ice-water bath. After this, the homogenate was centrifuged at 570 g for 10 min. The supernatant was collected. After this, the content of triglyceride was analysed in accordance with the manufacturer’s instruction.

### Western Blot

Western blotting was used to detected the protein expression. The liver tissues were homogenized with RIPA containing a phosphate inhibitor cocktail for 30 min on ice and then centrifuged at 20,000 g for 15 min in 4 °C. The supernatant was collected and Protein concentration was determined with the commercial BCA Protein Assay Kit. Proteins were denatured by boiling at 95 °C for 5 min with loading buffer. Then 10% sodium dodecyl sulfate polyacrylamide gel electrophoresis (SDS-PAGE) were used to separate the denatured proteins. After that, protein extracts were transferred onto polyvinylidene fluoride (PVDF) membranes. Then, Tris Buffered Saline with Tween (TBST) containing 5% BSA was used to block the nonspecific binding site for 2 h at room temperature and the PVDF membranes were incubated overnight at 4 °C with primary antibodies. After incubation, membranes were then washed three times for 10 min each with TBST buffer. HRP-conjugated goat anti-rabbit secondary antibody (1:3000 diluted in TBST) were then applied for 1 h at room temperature. The protein bands were visualized by an ECL chemiluminescence reagents following the guidelines and semi-quantitative analysis was performed using Quantity One Software. β-Actin was used as an internal control for normalization.

### Statistical Analysis

All data were expressed as mean ± standard deviation (SD) and one-way ANOVA with LSD *t* test were used to determine the difference between two groups. Statistical analysis was performed with Graph Pad Prism 5 and *p* < 0.05 was considered statistically significant.

## Results

### PEMF Protected the db/db Mice from Oxidative Stress Injury

To investigate whether PEMF exposure could decrease liver oxidative stress in db/db mice, the GSH and GSSG level as well as MDA level, and antioxidant enzymes’ activity and expression were detected. As is shown in Fig. 2A-C, PEMF exposure significantly increased the reduced GSH level and decreased the MDA and GSSG level, comparing with sham group. Meanwhile, the activity of several antioxidant enzymes was tested including CAT, SOD and GSH-Px as is shown in Fig. 2D-F. The results indicated that PEMF exposure promoted the GSH-Px level except CAT and SOD level. In Fig. 2G, the expressions of CAT, SOD1 and SOD2 proteins were not altered by PEMF. The trend of glutathione reductase (GR) was higher in comparison with sham group, but the difference was not statistically significant. Only the overexpression of GSH-Px was observed significantly.

**Figure 2:**
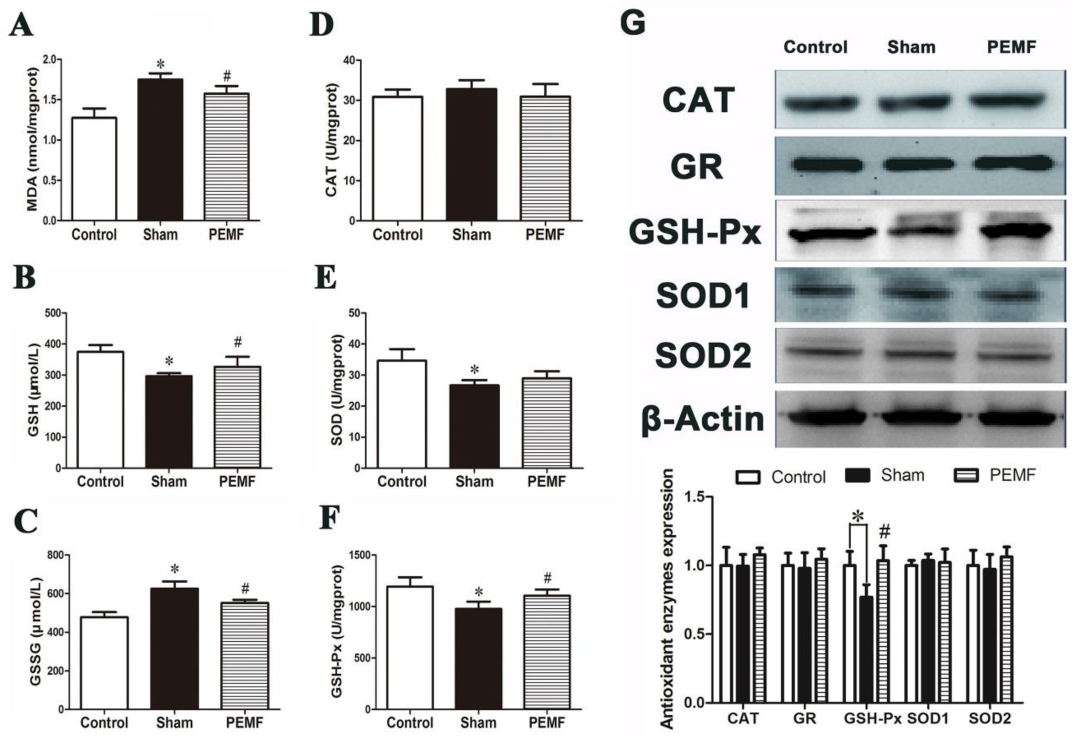
Effects of PEMF exposure on redox homeostasis of db/db mice. **A** Liver MDA content. **B** Liver GSH content. **C** Liver GSSG content. **D** Activity/unit of CAT. **E** Activity/unit of SOD. **F** Activity/unit of GSH-Px. **G** Expression of CAT, GR, GSH-Px, SOD1 and SOD2 were determined through Western Blotting. Values are all expressed as mean ± SD. ^*^ *p*<0.05, compared with control; ^#^ *p*<0.05, compared with sham.

### PEMF Alleviated Hepatic Lipid Accumulation by inhibiting SREBP-1c Expression in db/db Mice

Fig. 3A shows that the liver weight was significantly decreased in PEMF group. Besides, PEMF exposure could reduce liver TG content by around 38 % (Fig 3B). Hepatic lipid accumulation was investigated by Oil Red O staining. As observed in Fig. 3C, both the number and size of lipid droplets were decreased remarkably after given PEMF stimulation. The expression of SREBP-1c was also attenuated significantly after given PEMF exposure as shown in Fig. 3D. These indicated that PEMF-treat could improve liver lipid metabolism.

**Figure 3:**
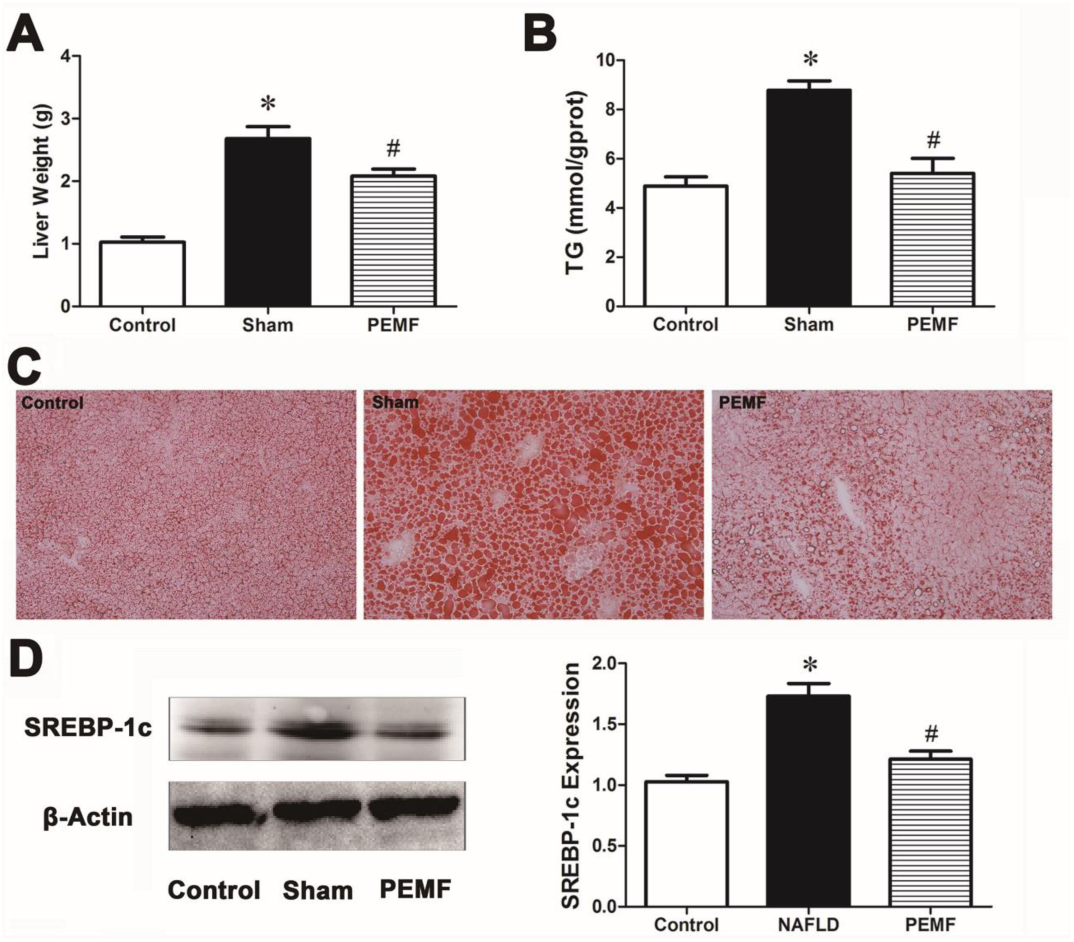
The liver change and SREBP-1c expression of mice. **A** Weight of liver. **B** Liver triglyceride content was measured with commercial available kits. **C** Liver lipid accumulation was evaluated by Oil Red O staining (×200). **D** Liver SREBP-1c expression was determined by Western Blotting. Values are all expressed as mean ± SD. ^*^ *p*<0.05, compared with control; ^#^ *p*<0.05, compared with sham.

## Discussion

NAFLD has become one of the most liver disease worldwide and affects people quality of life. And the continuously increasing trend of NAFLD with the prevalence and incidence of T2DM is alarming^18, 19^. To date, the pathogenesis of NAFLD has not been completely clarified, and there is no accepted treatment available for NAFLD^1, 5^. Oxidative stress has been recognized as a key mechanism affect the progression of NAFLD in T2DM, by inducing severe variations in lipid metabolism^20-22^. Therefore, antioxidant therapeutic approach has become an attractive treatment for diabetic hepatic complications. Accordingly, the purpose of this present study was to investigate whether PEMF exposure could alleviate oxidative stress and lipids accumulation in diabetic liver to improving NAFLD. The main finding of our present study was that PEMF exposure reduced oxidative stress in association with attenuated hepatic steatosis in db/db mice.

Nowadays, there are evidences suggesting that ROS plays the causal role in liver lipid droplets formation, while some others with the results that abnormal accumulation of lipid in liver induced ROS overproduced^23, 24^. Thus, ROS serves as a key driving force to exacerbates the progression from initial hepatic steatosis to the further NAFLD. It was already reported that antioxidant molecules were depleted and antioxidant enzymes were inactivated by ROS^25^. In this study, compared with sham group, the content of MDA in PEMF group, as an indicator of oxidative stress, was significantly reduced. While GSSG level was obviously decreased in PEMF group compared with sham group. Meanwhile, GSH level was increased. MDA is a major end-product, thus as an index of lipid peroxidation. And it cross-links with protein and nucleotides^26^. Up to this, our results showed that redox homeostasis was improved by given PEMF exposure.

To further explore the mechanisms that how the PEMF exposure regulated the oxidative stress, we investigated the activity and expression of a series of antioxidant enzymes. The activity/unit and expression of CAT were not changed as the same as SOD. Expression of GR had increasing trend but was not statistically significant. Both of the activity/unit and expression of GSH-Px were significantly increased. For its abundance, GSH serves as a cellular reductant and metabolic controller which is able to scavenge the free radicals against oxidative damage^27^. In order to achieve the purpose of counteract peroxide, the GSH-Px would promote the reaction between GSH and H_2_O_2_^28^. And it could convert the GSH (reduced form of glutathione) as a co-substrate into GSSG (oxidized form of glutathione). GSH-Px as enzymes could scavenge the hydrogen peroxide and lipid hydroperoxides so as to prevent the peroxides damage^26, 29^. Afterwards, GSSG converts into GSH catalysed by GR. When the level of GSH decreased, the antioxidant status got diminished and this would lead to the enhanced lipid peroxidation, even worse, indicate the apoptotic state of cells^30^. Recent study reported that, including 879 cases of T2DM and 1295 healthy controls, the GSH-Px activity was decreased obviously, while the plasma MDA concentration was increased significantly in insulin resistance individuals in comparison of controls^31^. As newly reported, acupuncture could inhibit oxidative stress with increasing activity of GSH-Px in abdominal obese NAFLD rats^32^. Presumably, PEMF exposure could ameliorate hepatic oxidative stress in db/db mice by increasing the activity/unit and expression of GSH-Px and then enhance the antioxidant ability of enzymes.

Excess accumulation of triglyceride in the hepatocyte is another characteristic of NAFLD. Until now, previous studies already have implied that *de novo* lipogenesis leads considerably to redundant hepatic lipid storage and steatosis in patients with NAFLD^33, 34^. It is traditionally thought that abnormal lipid accumulation induced mitochondrial dysfunction and the mitochondrial stress generated ROS^34, 35^. To explore the beneficial effect of PEMF on NAFLD, this study also payed attention to the change of liver which is the most important organ in body metabolism. After 12-week administration, the liver weight, TG content determination and frozen sections proved that PEMF exposure could obviously decreased the hepatic weight through scavenging lipid droplets in hepatocytes compared with sham group. Some findings suggested that electromagnetic fields exposure could induce lipid profile changes on brain and is similar to physiological stress^36^. There are also some results indicated that electromagnetic fields exposure could induce effects on soil nematodes through alternation of lipid metabolism^37^. The present study had the similar findings with that electromagnetic fields exposure could induce effects on lipid metabolism.

To investigate the potential key transcription factor, this study focused on sterol regulatory element binding proteins (SREBPs) that regulate the expression of genes control synthesis of fatty acid and cholesterol. Previous studies have identified there are three subtypes in SREBPs: SREBP-1a, SREBP-1c and SREBP2. SREBP-1c is expressed with specially high level in the fatty livers of obese, insulin resistant and hyperinsulinaemic animal models^38, 39^. Since PEMF could significantly improve the lipid accumulation in db/db mice, this study investigated the protein expression level of SREBP-1c. And the result showed that PEMF exposure attenuated the expression of SREBP-1c by about 30%. Huang et al reported that Meretrix meretrix oligopeptides downregulated SREBP-1c expression to ameliorate high-fat-diet-induced NAFLD^40^. Xyloketal B had an effect of lowering hepatic lipid accumulation via targeting the activation of SREBP-1c on NAFLD^20^. Hepatic lipid accumulation also could be attenuated by Crude triterpenoid saponins from ilex latifolia via inhibiting SREBP-1c expression on NAFLD^41^. Therefore, the present study indicated that PEMF exposure could reduce abnormal liver lipid accumulation via downregulating the SREBP-1c expression in db/db mice.

## Conclusion

In conclusion, PEMF exposure could protect liver from oxidative injury by increasing antioxidant enzyme activity and eliminate lipid accumulation by declining the SREBP-1c expression. Although the exact pathological process of NAFLD has been unclear and there is no specific therapeutic strategies with NAFLD, the present study probably provides a new solutions for NAFLD prevention and treatment.

## Acknowledgements

This work was supported by the Natural Science Foundation of Shaanxi Province (2018SF176) and National Nature Scientific Foundation (NSFC81471806 and NSFC51577188).

